# Increases in *BCL2L1* and *ID1* dosage synergistically drive fate bias and competitive advantage in human pluripotent stem cells

**DOI:** 10.64898/2026.03.26.714405

**Authors:** Yingnan Lei, Nuša Krivec, Arkajyoti Sarkar, Mai Chi Duong, Anfien Huyghebaert, Charlotte Janssens, Stefaan Verhulst, Leo A. van Grunsven, Diana Al Delbany, Claudia Spits

## Abstract

**Background:** Gains of chromosome 20q11.21 are among the most common culture-acquired abnormalities in human pluripotent stem cells (hPSC), conferring a well-defined survival advantage while altering differentiation capacity. However, it remains unclear whether this advantage persists during differentiation, how the aneuploidy alters ectodermal and retinal pigment epithelium (RPE) lineage specification, and which genes within the minimal amplicon drive these effects.

**Methods:** We used three isogenic human embryonic stem cell line pairs (wild-type and 20q11.21 gain) and assessed their behaviour in two neuroectoderm differentiation systems: directed neuroectoderm induction (dual SMAD inhibition) and long-term spontaneous RPE differentiation. Competitive dynamics were measured in mixed cultures, and lineage outcomes were analysed using immunostaining, gene expression profiling and single-cell RNA sequencing. To identify driver genes, we generated *BCL2L1* and *ID1* overexpression lines and tested their effects under both directed and spontaneous differentiation conditions.

**Results:** Across all lines and conditions, 20q cells expanded from a minor fraction to dominate mixed cultures, indicating that their competitive advantage persists beyond the undifferentiated state. Despite this dominance, pure 20q cells failed to specify to neuroectoderm or RPE. Single-cell transcriptomics revealed consistent diversion toward non-neural ectodermal and extraembryonic fates. Mechanistically, overexpression of *BCL2L1* and *ID1* alone or in combination impaired neuroectoderm specification, while synergistic effect of both genes promoted non-neural ectodermal outcomes under directed differentiation conditions. In spontaneous differentiation, both genes could disrupt differentiation.

**Conclusions:** The 20q11.21 gain couples a persistent survival advantage with a disruption of neural and RPE lineage competence, redirecting cells toward alternative ectodermal and extraembryonic fates. These effects arise from the combined action of two dosage-sensitive genes *BCL2L1* and *ID1* within the amplicon, illustrating how regional gene dosage can reshape developmental signalling responses in hPSC.

## INTRODUCTION

As of March 2026, more than 8.800 human pluripotent stem cell (hPSC) lines have been registered in the European Human Embryonic Stem Cell Registry (hpscreg.eu), including 1.123 human embryonic stem cell (hESC) lines and 7.722 induced pluripotent stem cell (hiPSC) lines. To date, these lines have contributed to 205 clinical studies, predominantly targeting degenerative disorders of the central nervous system and ocular diseases, with clinical translation continuing to expand steadily [1–3]. The emergence of hPSC-technologies has addressed key limitations of traditional animal models, revolutionizing disease research by enabling the differentiation of patient-derived or genetically modified hiPSC/hESC into clinically relevant cell types [4–6].

Although hPSC hold substantial therapeutic and research potential, prolonged *in vitro* culture poses important challenges. Continuous adaptation to culture conditions can drive genetic and epigenetic alterations that undermine genomic stability and compromise cell line integrity. Recurrent chromosomal abnormalities are particularly well documented, most commonly gains of chromosomes 1, 12, 17 and 20, while deletions affecting regions such as 10p and 18q occur less frequently [7,8]. Point mutations have also been identified in cancer-related genes including *TP53* and *BCOR* [9–13], as well as mitochondrial DNA mutations [14] and epigenetic changes [15,16]

These abnormalities can disrupt normal self-renewal control and apoptotic responses, conferring a selective growth advantage that enables genetically altered clones to progressively outcompete and overtake normal cells in culture [8,16–18]. There is increasing evidence that genetic aberrations can significantly compromise the differentiation capacity of hPSC. For instance, hPSC with a loss of 18q or gains of 1q exhibit reduced neuroectoderm potential [18–20], trisomy 12 results in reduced spontaneous differentiation efficiency [21], and trisomy 17 drives aberrant lineage specification in embryoid bodies (EBs) [22]. Beyond lineage bias, several abnormalities have been linked to increased tumorigenic risk. hPSC carrying trisomy 12 have been reported to generate malignant teratocarcinoma-like tumours [23], while the presence of more than three copy number variants (CNVs) in iPSC correlates with abnormal tissues formation upon transplantation in mice [24]. In addition, aneuploidy in mouse ESC has been shown to enhance teratoma metastasis [25]. The significant alterations imposed by aneuploidy on the cellular behaviour raises important concerns regarding the safety and efficacy of stem-cell-based interventions and can have important consequences for their reliability as research models (Andrews et al., 2022; Benvenisty et al., 2025). For example, a clinical trial involving hiPSC-derived retinal pigment epithelium (RPE) was temporarily halted due to the detection of potentially deleterious mutations in one of the cell preparations intended for transplantation [27].

One of the most frequently recurring abnormalities in hPSC is duplication of chromosome 20q11.21, detected in more than 20 % of long-term cultures [7,11]. This same alteration is observed in several human cancers, where it has been linked to chemotherapy resistance and enhanced metastatic potential [28]. The minimal 20q11.21 amplicon spans 13 genes, several of which are associated to oncogenesis ([28]. Among them, *BCL2L1*, encoding the anti-apoptotic protein BCL-xL, is identified as a principal driver of the selective growth advantage associated with this gain, enhancing cell survival and reducing apoptosis under stress [29,30]. Our group previously showed that 20q11.21 gain impairs neuroectoderm differentiation and that *BCL2L1-*overexpression recapitulates this phenotype [31]. Independent studies have similarly reported an altered lineage balance in EBs bearing the 20q11.21 gain (Jo et al., 2020), and isochromosome 20q variants exhibit aberrant cell-fate outcomes during spontaneous RPE differentiation [33]. Conversely, more recent findings published during the course of this study indicates that hPSC carrying 20q11.21 gain can differentiate efficiently into RPE [34].

Despite growing insight into the biological impact of 20q11.21 gain, several fundamental questions remain unresolved, which we systematically address in this study. First, do hPSC harboring this gain retain their selective advantage during differentiation? If this advantage persists during differentiation, low-level mosaicism that escapes routine detection could drive genetic drift during lineage commitment, ultimately yielding heterogeneous and potentially unstable cell products. Second, when challenged with neuroectodermal induction, which alternative lineage trajectories are preferentially adopted by 20q11.21 cells? Third, is the neuroectodermal defect attributable solely to increased *BCL2L1* dosage or do additional genes within the 20q11.21 amplicon that cooperate to redirect cell-fate specification?

## RESULTS

### hESC^20q^ retain their competitive growth advantage during ectodermal differentiation

To determine whether the fitness advantage associated with 20q11.21 gain is maintained during ectodermal differentiation, we carried out co-culture competition assays in two models: an 8-day directed neuroectoderm induction and a long-term RPE spontaneous differentiation protocol. Three in-house hESC lines (VUB02, VUB03, VUB14) that had spontaneously acquired a 20q11.21 amplification during routine culture (Fig. S1 and Table S1) were fluorescently tagged (VUB02^20q-Venus^, VUB03^20q-Venus^, VUB14^20q-mCherry^). Each 20q variant line was mixed with its isogenic wild-type (WT) counterpart prior to differentiation, at an initial proportion of 10% variant to 90% WT in the neuroectoderm model and 5% variant to 95% WT in the RPE model. The relative proportion of 20q11.21-positive cells was quantified at defined timepoints with parallel time-lapse confocal imaging to monitor population dynamics.

By day 8 of directed neuroectoderm differentiation, 20q11.21-positive cells had expanded significantly in all three lines. The mutant fraction increased from 16.3 ± 0.9 % to 71.1 ± 2.4 % in VUB02 (p = 0.0001), from 17.2 ± 1.1 % to 43.2 ± 1.7 % in VUB03 (p = 0.0016), and from 11.7 ± 0.8 % to 64.0 ± 3.1 % in VUB14 (p = 0.0001). (Fig. 1A)

**Figure 1.**
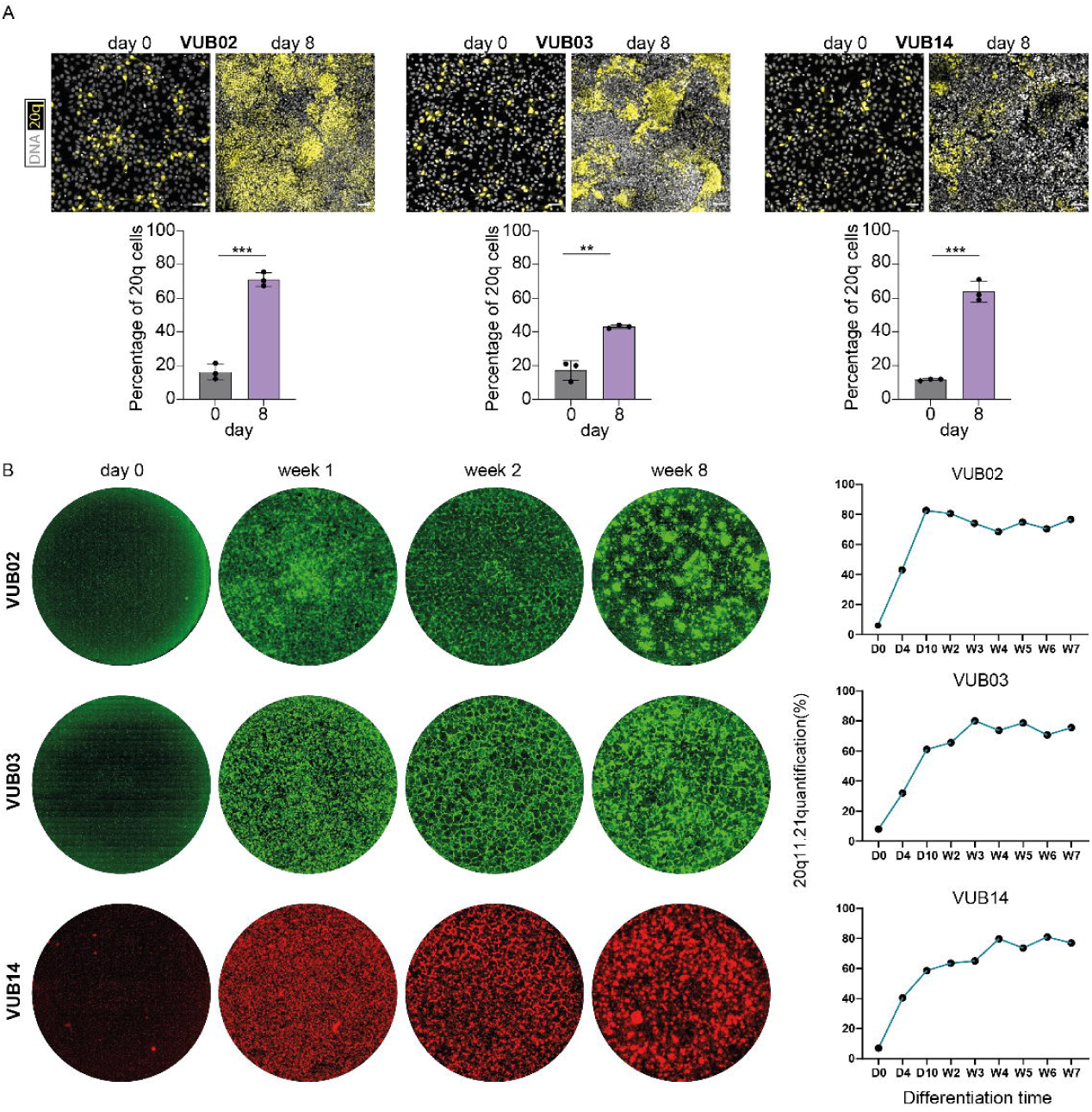
Competitive advantage of hESC^20q^ during directed and spontaneous neuroectodermal differentiation. (A) Representative images of co-culture conditions at initiation (day 0) and after directed neuroectoderm differentiation (day 8). hESC^20q^ (yellow, fluorescently labelled) were mixed with hESC^WT^ at a 1:9 ratio, and populations dynamics were tracked via fluorophore expression. The bar plots show quantification of 20q-positive cells at day 0 and day 8, measured using a cell counter (N = 3). (B) Representative images of co-cultures containing 5 % fluorescently labelled 20q cells and their isogenic balanced counterparts at day 0 and at weeks 1, 2, and 8 of spontaneous RPE differentiation. Line plots show the proportions of 20q-positive cells throughout differentiation, quantified by cell counting at early time points (D0 - D10, N = 2) and by weekly image-based analysis at later stages (W2-W7). * p<0.05, ** p<0.01, *** p<0.001, **** p<0.0001, ns=non-significant; unpaired t-test

A comparable dynamic was observed during spontaneous RPE specification. The fluorescent fraction increased rapidly during the first ten days and subsequently reached a plateau over the following seven weeks, with approximately 80 % of cells remaining fluorescent. Cell numbers were quantified until day 10, after which whole-well confocal imaging was used from week two to week seven. In VUB02, the proportion of 20q11.21-positive cells increased from 6.0 % to 82.5 % within the first ten days (p = 0.0012) and then remained relatively stable, fluctuating between 68.4 % and 80.5 %. A comparable pattern was observed in the two other lines, VUB03 and VUB14, where mutant fractions increased from 8.0% to 61.0% (p = 0.0014) and from 7.0% to 58.5% (p = 0.0005), respectively, before stabilizing between approximately 65% and 80% up to week seven (Fig. 1B).

Collectively, these results show that 20q11.21 duplication confers a persistent fitness advantage, enabling aneuploid cells to progressively overtake WT counterparts during both short-term neuroectoderm induction and long-term RPE differentiation. Importantly, this clonal dominance emerged in the absence of passaging, highlighting the intrinsic strength of the selective advantage exerted during differentiation itself rather than during routine expansion.

### 20q11.21 gain impairs directed and spontaneous neuroectodermal differentiation in hESC

We next assessed the impact of 20q11.21 gain on directed and spontaneous neuroectodermal differentiation by comparing three hESC^20q^ lines with their isogenic diploid counterparts. BH VDifferentiation efficiency was assessed by quantifying lineage-specific markers (SOX1, PAX6, BEST1, MITF, RPE65) at both the protein and transcript levels, while persistence of undifferentiated cells was evaluated using the pluripotency-associated marker OCT4.

Immunofluorescence analysis revealed a markedly reduced population of PAX6^+^ cells in all three hESC^20q^ lines after 8 days of induced differentiation compared to WT lines (Fig. 2A). Consistently, *PAX6* and *SOX1* mRNA levels were significantly lower in 20q cultures (p = 0.0005 and p = 0.0013, respectively; Fig. 2B), indicating reduced neuroectodermal differentiation efficiency. All differentiated populations showed minimal *OCT4* expression at the transcript level (Fig. 2B), and no detectable OCT4^+^ cells by immunofluorescence (Fig. 2A), confirming efficient exit from the pluripotent state in both WT and 20q lines.

**Figure 2.**
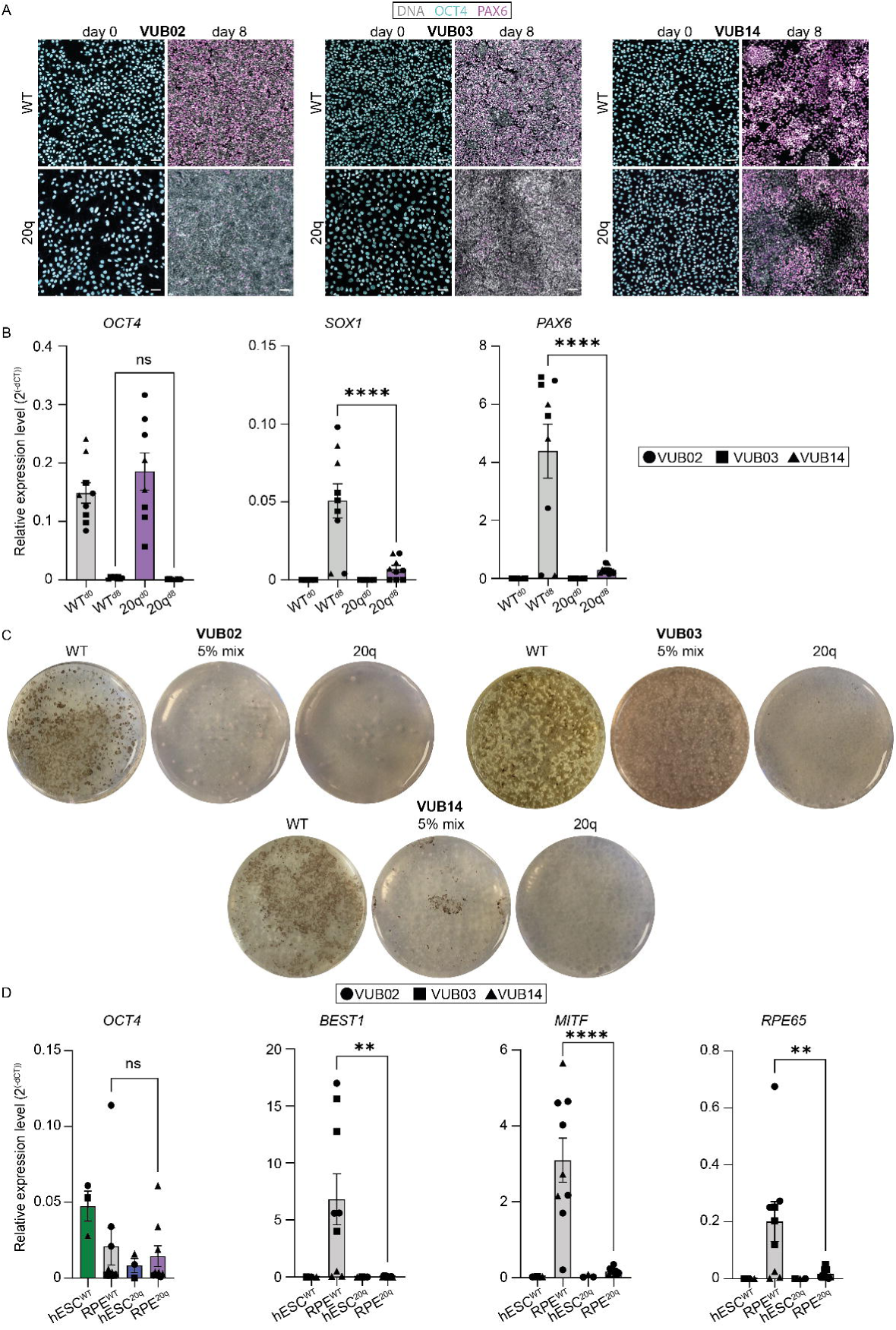
hESC^20q^ show impaired directed and spontaneous neuroectodermal differentiation (A) Immunostaining for OCT4 and PAX6 at day 0 and day 8 of directed neuroectoderm differentiation. Representative images are shown for three hESC^WT^ lines (VUB02^WT^, VUB03^WT^, VUB14^WT^) and three hESC^20q^ (VUB02^20q^, VUB03^20q^, VUB14^20q^). (B) Relative mRNA expression of *OCT4*, *SOX1*, and *PAX6* at day 0 and day 8 of neuroectoderm differentiation in three biological replicates per genotype. (C) Whole-well images after 3 months of RPE differentiation for hESC^WT^, 5 % hESC^20q^ co-cultures, and pure hESC^20q^ cultures. Successful RPE induction is characterized by the formation of pigmented colonies, as observed in WT cultures. **(D)** Relative mRNA expression levels of RPE markers (*BEST1*, *MITF, RPE65*) and the pluripotency marker *OCT4* in the differentiated WT and 20q cell lines, as well as in undifferentiated controls (N = 3-9 per line). * p<0.05, ** p<0.01, *** p<0.001, **** p<0.0001, ns=non-significant; unpaired t-test

Furthermore, following 3 months of RPE differentiation, cultures derived from hESC^WT^ were densely populated with pigmented cells, indicative of successful RPE formation (Fig. 2C). In contrast, hESC^20q^ cultures showed rare and scattered pigmented regions, suggesting a markedly reduced capacity to generate RPE (Fig. 2C). These morphological differences were confirmed by qPCR analysis for key RPE markers *BEST1, MITF* and *RPE65*, which were robustly induced in hESC^WT^ but remained low or undetectable in 20q samples (Fig. 2D; p = 0.0080, 0.0001, 0.0183, respectively). Notably, VUB03^20q^ and VUB14^20q^ retained elevated expression of *OCT4* (Fig. 2D), indicating the persistence of residual undifferentiated cells. Taken together, these findings show that hESC^20q^ exhibit impaired differentiation toward both neuroectodermal and RPE lineages, with persistent retention of pluripotent population even after prolonged spontaneous differentiation.

### 20q11.21 gain biases hESC differentiation toward undifferentiated and amnion or surface ectoderm identities

To characterize the cellular identities underlying the altered differentiation behaviour of 20q cells, we performed single-cell RNA sequencing (scRNA-seq) on hESC^WT^ and hESC^20q^ following 8 days of directed neuroectoderm differentiation and 60 days of spontaneous differentiation.

UMAP visualization of cells following directed neuroectoderm differentiation revealed clearly segregated transcriptional landscapes between hESC^WT^ and hESC^20q^ derivatives (Fig. 3A). Cell clusters were annotated based on UCell-derived lineage signature scoring using established marker gene sets (Fig. S2), and the corresponding lineage enrichment scores across clusters are shown in Fig. 3B. We next quantified the proportions of annotated cell types and found that more than 90% of hESC^WT^-derived cells acquired the expected neuroectodermal identity, with a minor fraction progressing toward more mature neuronal states (Fig 3C). In contrast, hESC^20q^-derived cells exhibit a substantially reduced neuroectoderm fraction (0–46%; Fig. 3C), indicative of lineage diversion toward alternative lineage identities, with line-specific differentiation trajectories. VUB03^20q^ predominantly adopted a surface ectoderm identity, with a smaller subset expressing anterior neuroectoderm markers OTX2 or SIX3 (Fig. 3C). VUB14^20q^ exhibited a more balanced neuro-surface-ectoderm composition and induced a minor anterior neuroectoderm population. VUB02^20q^ was largely characterized by a separate cluster enriched for early neuroectoderm-associated transcripts but lacking canonical neuroepithelial markers such as PAX6 (Supplementary Data File 1), consistent with a transcriptionally divergent, non-canonical neuroectodermal state. This line additionally comprised surface ectoderm cells and a unique proliferative cluster (Fig. 3C). To validate these findings, we carried out independent replicates of directed neuroectoderm differentiation and stained them for neuroectoderm (PAX6) and surface ectoderm markers (KRT8, TFAP2A, EPCAM), as well as OCT4 (Fig. 3D, 3E and S3A). The 20q cultures showed significantly reduced fractions of PAX6^+^ cells as compared to WT cells (6.2 % vs 81.8 %, p = 0.0028), together with increased proportions KRT8^+^ (52.5 % vs 9.5 %, p = 0.0579), TFAP2A^+^ (11.1 % vs 0 %, p = 0.0813) and EPCAM^+^ cells (83.5 % vs 16.0 %, p = 0.0026). OCT4 was undetectable in all differentiated cultures, confirming complete exit from pluripotency.

**Figure 3.**
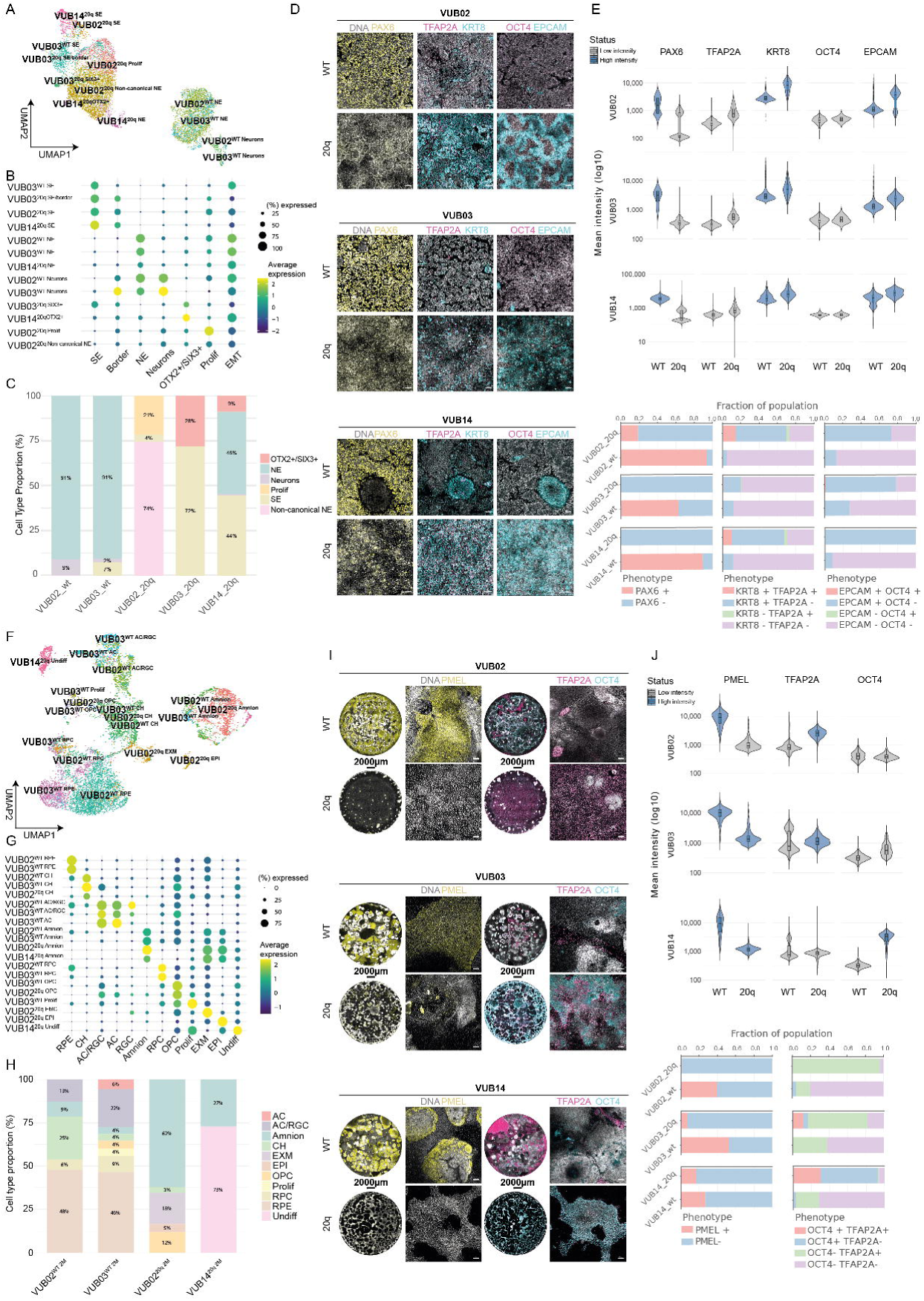
Cell fate specification during directed (panels A-E) and spontaneous (panels F-J) neuroectoderm differentiation. **(A)** UMAP visualization of two hESC^WT^ lines (VUB02 and VUB03) and three hESC^20q^ lines (VUB02, VUB03, and VUB14). Neuroectoderm (NE), neurons, surface ectoderm (SE), surface ectoderm with border genes co-expression (SE/border), proliferating cells (Prolif), anterior ectoderm (OTX2+/ SIX3+), non-canonical neuroectoderm (Non-canonical NE). **(B)** UCell scoring of ectodermal lineage signatures across the clusters shown in (A). Bubble size: proportion of cells expressing lineage genes. Color: average expression level. **(C)** Relative proportions of each identified cell type across the individual cell lines. **(D)** Immunofluorescence staining: PAX6 (yellow, neuroectoderm; panel 1), TFAP2A (magenta, surface ectoderm; panel 2), KRT8 (turquoise, surface ectoderm; panel 2), OCT4 (magenta, pluripotency; panel 3) and EPCAM (turquoise, surface ectoderm; panel 3). **(E)** Quantification of the neuroectoderm immunostaining showing marker intensity and the fraction of marker-positive cells relative to the nuclear stain. Intensity was measured as mean fluorescence intensity, and marker-positive cells were defined based on a predefined threshold (unpaired t-test, N^wt^=3, N^20q^=3). **(F)** UMAP visualization of WT (VUB02, VUB03); 20q11.21 duplication (VUB02 and VUB14). Cortical hem (CH), amacrine cells or retinal ganglion cells (AC/RGC), retinal progenitor cells (RPC), retinal pigment epithelium (RPE), oligodendrocyte progenitor cells (OPC), proliferating (Prolif), extra-embryonic mesoderm cells (EXM), epithelial cells (EPI), undifferentiated cells (Undiff). **(G)** UCell scoring of ectodermal lineage signatures across the clusters shown in (F). Bubble size: proportion of cells expressing lineage genes. Color: average expression level. **(H)** Relative proportions of each identified cell type across the individual cell lines. **(I)** Immunofluorescence staining after 2 months of spontaneous differentiation for key lineage markers: PMEL (yellow, RPE), TFAP2A (magenta, amnion), and OCT4 (turquoise, pluripotency). **(J)** Quantification of the RPE immunostaining showing marker intensity and the fraction of marker-positive cells relative to the nuclear stain. (unpaired t-test, N^wt^=3, N^20q^=3).

Similarly, the scRNA-seq results for the spontaneous differentiation revealed genotype and cell-line dependent clustering (Fig. 3F). Cell lineage gene-expression signature analysis showed that spontaneous differentiation yields an array of cell identities (Fig. 3G). After two months of differentiation (Fig. 3H), approximately half of hESC^WT^ cells acquired RPE identity, with the remaining cells distributed among oligodendrocyte progenitor, retinal progenitor, amnion, amacrine/retinal ganglion, and cortical hem populations. In contrast, hESC^20q^ lines showed a complete absence of RPE specification and instead exhibited a strong bias toward extraembryonic lineages and residual undifferentiated cells. VUB02^20q^-cultures consisted mainly of amnion cells (62%) and extra-embryonic mesoderm cells (18%), while VUB14^20q^-cultures were largely composed of undifferentiated cells (73%) with the remaining cells adopting an amnion identity (27%). In independent confirmatory differentiation experiments, all 20q lines showed reduced proportions of PMEL⁺ cells (8.5% vs 39.8%, *p* = 0.0233) and increased TFAP2A⁺ populations (68.2% vs 26.4%, *p* = 0.1080) compared with WT (Fig. 3I–J). Consistent with the scRNA-seq results, VUB02^20q^ cultures were primarily TFAP2A⁺ (94.3%). In contrast, VUB14^20q^ retained OCT4 expression in 93.1% of cells, indicating a large fraction of residual undifferentiated hESCs. VUB03^20q^ displayed two distinct OCT4⁺ and TFAP2A⁺ populations, reflecting a mixture of residual undifferentiated hESCs and amnion cells (OCT4⁺ = 19.9%, TFAP2A⁺ = 74.2%) (Fig. 3I–J).

Taken together, these results indicate that duplication of 20q11.21 alters lineage commitment in hESCs, leading to distinct differentiation outcomes compared to WT cells depending on the differentiation conditions. During directed neuroectoderm differentiation using dual SMAD inhibition, 20q11.21 gain redirects differentiation away from the expected posterior neuroectoderm [35,36] toward heterogeneous ectodermal identities, including non-canonical ectodermal progenitors and anterior neuroectoderm, with surface ectoderm emerging as a recurrent lineage outcome across all lines. Moreover, under spontaneous differentiation conditions that favour a more anterior ectodermal environment leading to RPE formation [20], 20q cells predominantly adopt an amnion-like identity or remain undifferentiated.

### Overexpression of *BCL2L1* disrupts neuroectoderm differentiation and synergistically with *ID1* induces amnion and surface ectoderm misspecification

To identify candidate driver genes underlying the disruption of neuroectoderm differentiation observed in 20q cells, we investigated *BCL2L1* and *ID1*, two genes located within the minimal common region of the 20q11.21 gain and previously implicated in regulating cell survival and differentiation pathways relevant to pluripotency and lineage specification [7,30,31]. *BCL2L1* overexpression has been associated with deregulation of TGFβ/SMAD signalling, a pathway required for proper neuroectoderm differentiation [37,38], while *ID1*, a downstream target of BMP4 signalling [39,40], represents a plausible contributor to the observed differentiation bias. Therefore, we generated genetically balanced VUB02, VUB03, and VUB14 hESC lines overexpressing *BCL2L1* and *ID1* using lentiviral transduction, either individually or in combination. Figure 4A shows their expression in parental and transduced cell lines; relative expression of *BCL2L1* and *ID1* was lowest in control hESC^WT^, moderately increased in hESC^20q^, and highest in the genetically engineered lines.

**Figure 4.**
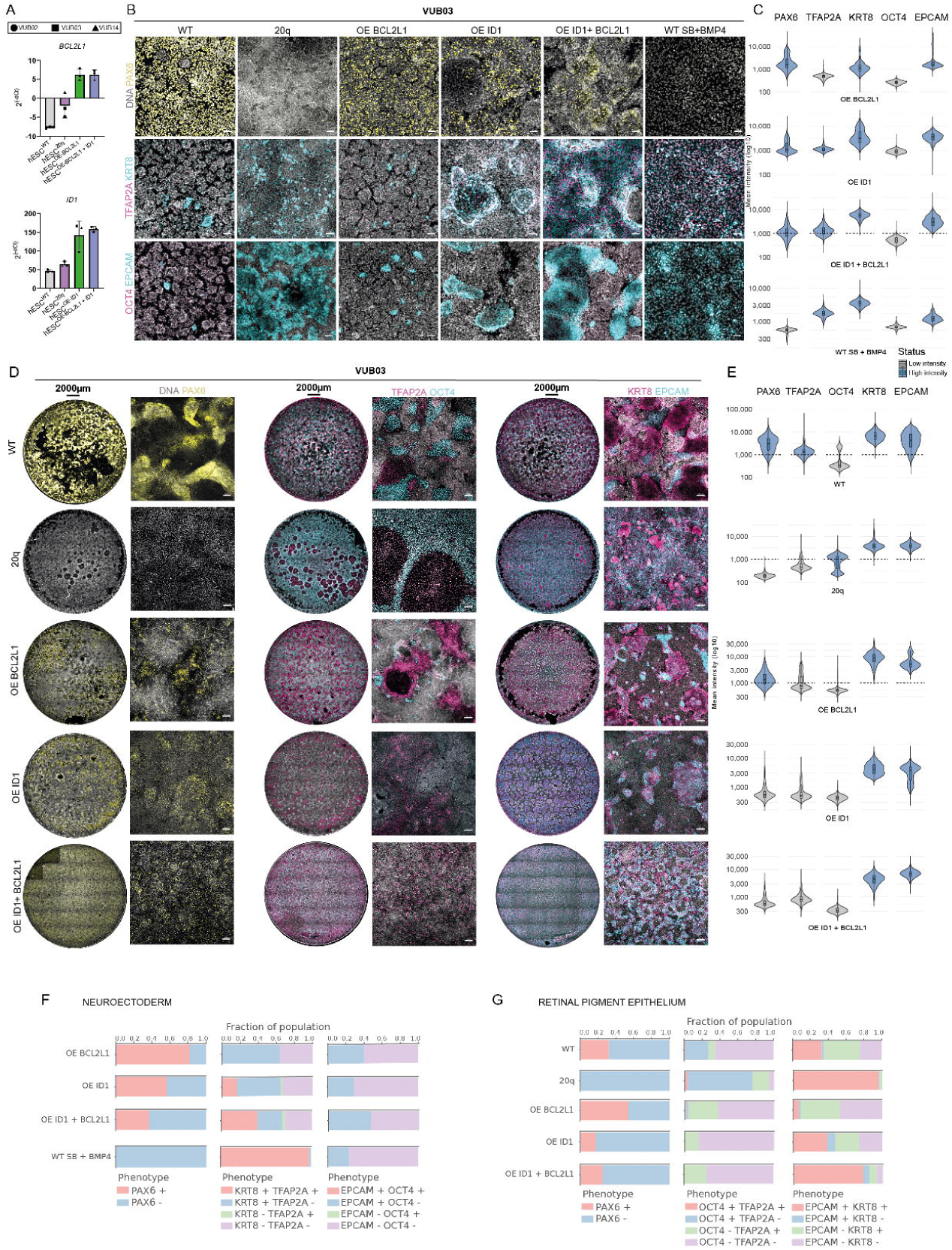
Effects of BCL2L1 and ID1 overexpression on hESC differentiation. **(B)** Quantitative RT-PCR analysis of BCL2L1 and ID1 expression levels in WT, 20q gain, and hESC lines overexpressing BCL2L1, ID1, or both (BCL2L1/ID1). **(C)** Immunofluorescence staining of the VUB03 cell line: WT, 20q gain, BCL2L1-OE, ID1-OE, and double (BCL2L1 + ID1) overexpression lines. The final condition represents wild-type cells treated with SB and BMP4. Stained markers include PAX6 (yellow; neuroectoderm, panel 1), TFAP2A (magenta; surface ectoderm, panel 2), KRT8 (turquoise; surface ectoderm, panel 2), EPCAM (turquoise; surface ectoderm, panel 3), and OCT4 (magenta; pluripotency, panel 3). **(D)** Quantification of neuroectoderm immunostaining of BCL2L1-OE, ID1-OE, and double (BCL2L1 + ID1) overexpression lines showing marker intensity similar to Fig. 3E. **(E)** Immunofluorescence staining of VUB03 cells after 14 days of spontaneous differentiation. Stained markers include PAX6 (yellow; neuroectoderm, panel 1), TFAP2A (magenta; surface ectoderm, panel 2), OCT4 (turquoise; pluripotency, panel 2), EPCAM (turquoise; surface ectoderm, panel 3), and KRT8 (magenta; surface ectoderm, panel 3). **(F)** Quantification of immunostaining of BCL2L1-OE, ID1-OE, and double (BCL2L1 + ID1) overexpression lines after spontaneous differentiation showing marker intensity similar to Fig. 4C. **(G)** Fraction of marker-positive cells after 8 days of neuroectoderm differentiation. **(H)** Fraction of marker-positive cells after 14 days of spontaneous differentiation.

Next, we subjected all lines to directed neuroectoderm differentiation for 8 days (Fig. 4B, 4C, 4F, S3A). Consistent with earlier results, WT and 20q lines showed the same differentiation phenotypes, with efficient PAX6⁺ neuroectoderm specification in WT cultures and reduced neuroectoderm commitment with increased surface ectoderm markers in 20q cells. *BCL2L1* overexpression *(BCL2L1*^OE^) alone did not disrupt neuroectoderm induction, with most cells remaining PAX6⁺ (87.6%). Although elevated *BCL2L1* increased the proportion of KRT8⁺ and EPCAM⁺ cells, TFAP2A expression remained minimal (67.0% KRT8⁺, 35.0% EPCAM⁺, 2.8% TFAP2A⁺), indicating that BCL2L1 alone did not fully recapitulate the misspecification phenotype observed in hESC^20q^ cells. *ID1* overexpression reduced the fraction of PAX6⁺ cells (56.4%) and increased the proportion of cells expressing all three surface ectoderm markers (64.8% KRT8⁺, 28.9% EPCAM⁺, 18.8% TFAP2A⁺). Notably, combined overexpression of *BCL2L1* and *ID1* produced the strongest reduction in neuroectoderm specification (37.0% PAX6⁺) and generated a population with marker expression patterns closely resembling those observed in hESC^20q^ cultures (67.3% KRT8⁺, 46.8% EPCAM⁺, 43.8% TFAP2A⁺). Given that surface ectoderm differentiation is promoted by high levels of BMP4 signaling [41], we tested whether treatment of hESC^WT^ with the TGFβ inhibitor SB431542 and the BMP4 agonist could recapitulate this misspecification path (Fig. 4B). Under these conditions, cells expressed high levels of TFAP2A, KRT8, and EPCAM (97.8%, 99.5%, and 23.4%, respectively), consistent with the surface ectoderm phenotype observed in *BCL2L1/ID1* co-overexpression and in 20q gain lines, suggesting that this alternative cell fate may arise through this route.

Next, we examined the differentiation behavior during 14-day spontaneous differentiation, corresponding to the initial phase of RPE differentiation (Fig. 4D, 4E, 4G, S3B, S3C and S4). At this stage of spontaneous differentiation, hESC^WT^ cultures showed on average 71% PAX6⁺ cells and contained smaller populations of surface ectoderm and undifferentiated cells (14.2% TFAP2A⁺, 61.1% KRT8⁺, 18.5% EPCAM⁺, 9.1% OCT4⁺). Similarly to the directed differentiation condition, hESC^20q^ cultures showed no detectable PAX6 expression and contained a substantial fraction of residual undifferentiated cells (45.1% OCT4⁺) and surface ectoderm cells (33.0% TFAP2A⁺, 94.2% KRT8⁺, 89.9% EPCAM⁺).

hESC lines overexpressing *BCL2L1*, *ID1*, or both showed reduced fractions of PAX6⁺ cells (37.0%, 17.5%, and 25.2%, respectively) together with increased proportions of TFAP2A⁺ (48.1%, 15.4%, 25.1%), KRT8⁺ (74.3%, 66.3%, 86.1%), and EPCAM⁺ cells (48.0%, 47.0%, 85.1%), consistent with a shift toward amnion-like identity compared with WT cultures. In contrast, residual pluripotency was observed only in BCL2L1^OE^ lines (21.8% OCT4⁺), whereas ID1^OE^ and BCL2L1/ID1^OE^ lines showed no detectable OCT4 expression. We additionally examined the trophoblast marker GATA3 and the extraembryonic mesoderm marker HAND1, which were expressed in 20q and *BCL2L1*^OE^ cells in a cell line–specific manner (Fig. S4).

Together, these results demonstrate that during directed differentiation, *BCL2L1* and *ID1* act synergistically to disrupt neuroectoderm specification and recapitulate the alternative surface ectoderm fate observed in 20q cells. In contrast, during spontaneous differentiation, overexpression of either *BCL2L1* or *ID1* alone is sufficient to perturb normal differentiation, leading to the induction of amnion-like identities and in the case of BCL2L1, the persistence of residual pluripotency.

## DISCUSSION

In this study, we investigated how the recurrent 20q11.21 gain influences lineage specification during neuroectoderm differentiation of hPSC using two protocols that generate neural progenitors and retinal pigment epithelium, cell types under active investigation for clinical applications, and aimed to identify the molecular drivers underlying these effects. While previous work established that the 20q11.21 gain, through overexpression of the anti-apoptotic gene *BCL2L1* (BCL-xL) confers a survival advantage in the undifferentiated state [29,30] and disrupts neuroectoderm formation while largely preserving mesendoderm potential [31], it remained unclear whether this defect reflects a general failure to differentiate or a redirection toward alternative developmental identities.

Our data show that the competitive advantage conferred by the 20q11.21 gain extends beyond the pluripotent state into ectodermal differentiation, enabling mutant cells to dominate mixed populations during both neuroectoderm induction and spontaneous RPE differentiation. By characterizing the differentiation trajectories of hPSC^20q^, we further show that neuroectoderm specification is not simply impaired but associated with the emergence of alternative cell fates. This work extends a line of research from our group aimed at understanding how recurrent hPSC aneuploidies influence competitive behaviour and lineage outcomes during differentiation. Consistently, we previously demonstrated that hESCs carrying a gain of 1q outcompete their isogenic WT counterparts not only in the undifferentiated state but also during differentiation toward neuroectoderm, hepatoblast, and cardiac progenitor lineages in both 2D monolayers and 3D embryoid bodies through elevated expression of *MDM4* [18]. More recently, we showed that spontaneous RPE differentiation acts as a strong bottleneck for mosaic aneuploidy: while most recurrent abnormalities—including trisomy 20, isochromosome 20q, 17q gains, and 18q losses—are progressively depleted and fail to generate correctly specified RPE, 1q-gain cells persist, expand, and contribute substantially to the final RPE monolayer when co-cultured with WT cells [42]. This underscores the importance of identifying misspecified identities to clarify how culture-adapted aneuploidies reshape developmental trajectories and generate unintended cell populations during directed differentiation of pluripotent stem cells.

During directed neuroectoderm differentiation, 20q cells were redirected toward non-neural ectodermal identities. A similar pattern was observed during spontaneous RPE differentiation, where hESC^20q^ again failed to adopt the neural/RPE lineages and instead generated mixtures of non-neural ectodermal and amnion-like populations, with residual undifferentiated cells persisting in some lines. Notably, surface ectoderm and amnion share transcriptional features of the non-neural ectoderm lineage, which emerges through a transitional state marked by increased surface ectoderm gene expression before activation of the amniotic ectoderm program, and where cell density plays an important role in determining cell fate [43,44].

Our findings align with previous studies showing that aneuploidy frequently impairs neuroectodermal differentiation and extend our earlier observations for 1q gain as well as reports on isochromosome 20q [18,19,31,33,42]. This convergence suggests a broader developmental response in which aneuploid cells are either eliminated or diverted toward peripheral lineages, such as surface ectoderm and amnion, where they may pose less risk to the developing embryo. This is in line with work on hPSC-based gastruloids, where aneuploid cells were preferentially allocated to the trophectodermal lineage [45].

To dissect the molecular basis of the 20q11.21-associated fate deviations, we focused on candidate genes within the minimal amplicon with established roles in cell survival and lineage regulation, identifying *BCL2L1* and *ID1* as key contributors to the altered differentiation phenotype. Functional overexpression experiments showed that expression of either gene alone reduced the proportion of PAX6⁺ neuroectodermal cells but did not fully alter lineage outcomes. In contrast, their combined overexpression was sufficient to disrupt neuroectoderm specification and reproduce the shift toward surface ectoderm–like identities observed in 20q cells. Consistent with this model, co-expression of both genes decreased the proportion of PAX6⁺ cells under dual SMAD inhibition, supporting their role in modulating cellular responsiveness to inhibitory TGF-β cues [31]. Mechanistically, ID1, a canonical BMP4 effector, may sustain BMP signaling despite pharmacological inhibition of BMP receptors, thereby biasing cells away from neural fates. Although the precise molecular link between BCL2L1 and TGF-β signaling remains unresolved, previous studies have implicated BCL2L1 in later stages of neural development, including synapse maturation [46–48], suggesting that its influence on neural identity may extend beyond early specification events.

Spontaneous differentiation revealed greater sensitivity to individual perturbations than the tightly controlled conditions of directed differentiation, as overexpression of either *BCL2L1* or *ID1* alone was sufficient to disrupt normal specification, leading to the emergence of amnion-like identities and the persistence of residual pluripotency. This highlights the distinct signalling landscape of spontaneous differentiation, where cells exit pluripotency without externally imposed lineage constraints and are therefore exposed to fluctuating endogenous cues. Given that BMP4 signals can promote either ectodermal or mesendodermal fates depending on context, the full 20q-associated fate bias likely arises from the combined effects of enhanced survival, altered responsiveness to inhibitory signals, and variable endogenous BMP/TGF-β activity.

A recent study reported increased RPE differentiation efficiency in hPSC carrying a 20q11.21 gain and identified BCL-xL as the driver of this effect [34]. This contrasts with our results, where 20q cells consistently exhibited impaired RPE differentiation. Several factors may account for this discrepancy. This previous study relied on sublines derived from a single hESC line and genomic characterisation methods with limited resolution (G-banding, FISH, qPCR), which may not detect smaller CNVs arising during clonal expansion that could influence differentiation behaviour. Moreover, the use of subclones from a single genetic background may have shaped the observed outcomes, as the impact of chromosomal abnormalities on differentiation can vary depending on the genomic context of the cell line. Further work will be necessary to clarify the basis of these divergent observations.

Taken together, our findings indicate that the phenotypes associated with the 20q11.21 gain cannot be attributed to a single dominant driver but instead emerge from the cooperative activity of multiple genes within the amplicon. In our system, *BCL2L1* and *ID1* act together to override neural-inducing cues and promote alternative ectodermal outcomes, while elevated expression of either gene alone is sufficient to destabilize normal lineage progression in less constrained environments.

In summary, this study supports a model in which the phenotypes associated with the 20q11.21 gain arise from the cooperative activity of multiple genes within the amplicon rather than from a single dominant driver, highlighting the importance of regional gene dosage in shaping hPSC. These observations underscore the need for rigorous genetic monitoring of hPSC cultures, as chromosomal imbalances can subtly alter differentiation trajectories and cellular behaviour, with important implications for the safe and reliable translation of hPSC-based systems to clinical applications.

## EXPERIMENTAL PROCEDURES

### Resource availability

Requests for resources should be directed to the corresponding author, Claudia Spits.

### Materials availability

All VUB stem cell lines in this study, including the genetically abnormal sublines and genetically modified lines, are available upon request and after signing a material transfer agreement.

### Data availability

Raw sequencing data of human samples is considered personal data by the General Data Protection Regulation of the European Union (Regulation (EU) 2016/679), because SNPs can be extracted from the reads, and cannot be publicly shared. The data can be obtained from the corresponding author upon reasonable request and after signing a Data Use Agreement. The sc-RNA data are provided in the supplementary material, which allow for downstream gene-expression analysis. The data supporting all figures in this paper can be found at the Open Science Framework repository (https://osf.io/8p9qv/). Code, configuration files, and trained models for image quantification are publicly available at https://github.com/LIVR-VUB/IA-20q-REGE-2D/tree/main and https://huggingface.co/LIVR-VUB/20q-REGE-2D.

### Ethics statement

For all parts of this study, the design and conduct complied with all relevant regulations regarding the use of human materials, and all were approved by the local ethical committee of the University Hospital UZ Brussel and the Vrije Universiteit Brussel (File number: B.U.N. 1432020000669).

### hESC maintenance, passaging and transgenic modification

Human embryonic stem cell lines were derived and characterized as previously described (Mateizel et al., 2006, 2010) and are registered in the EU hPSC registry (https://hpscreg.eu/). All lines were routinely tested for Mycoplasma contamination (protocol: https://osf.io/jcen3/). VUB02 acquired a 1.45 Mb gain on chromosome 20q (VUB02^20q^), VUB14 a 1.68 Mb gain (VUB14^20q^), and VUB03 a 4.15 Mb gain (VUB03^20q^; Table S1, Fig. S1).

Cryopreserved stocks were stored in 90% KnockOut Serum Replacement (Thermo Fisher Scientific) and 10% DMSO. hESC were cultured on dishes coated with 5 µg/mL human recombinant Laminin-521 (Biolamina®) in NutriStem™ hESC XF medium (Biological Industries) supplemented with 10 U/mL penicillin/streptomycin (Thermo Fisher Scientific) at 37°C and 5% CO₂, with daily medium changes. For passaging, cells were dissociated with TrypLE™ (Thermo Fisher Scientific; 10 min, 37°C), inactivated with medium, centrifuged (1000 rpm, 5 min), and replated at 1:20–1:50 split ratios in medium supplemented with 1:100 RevitaCell (Thermo Fisher) for 24 h post-passaging. Karyotypic stability was confirmed by shallow genome sequencing, and all the lines were maintained within 10 post-thaw passages.

Fluorescently labelled hESC^20q^ lines were generated by lentiviral transduction as previously described[18]. Lentiviral particles were produced in HEK293T cells by co-transfection of VSV.G (pMDG) and gag–pol (pCMVΔR8.9) plasmids together with LeGO-EF1a-V2-Puro (Venus) or LeGO-EF1a-C2-Puro (mCherry) vectors, kindly provided by Kristoffer Riecken [51]. Viral supernatants were collected 48 and 72 h post-transfection. HPSC were transduced at a density of 10,000 cells/cm² using lentivirus in NutriStem/complete medium supplemented with protamine sulfate. Successfully transduced cells were enriched by fluorescence-activated cell sorting (FACS).

### Transgenic overexpression of BCL2L1 and ID1

The human BCL2L1 (BCL-xL) gene was cloned into the lentiviral vector pCDH (LV500A-1; System Biosciences). BCL-xL expression was driven by the elongation factor 1 alpha (EF-1α) promoter, while expression of GFP and the puromycin resistance gene was controlled by the phosphoglycerate kinase I (PGK) promoter. The ID1 expression plasmid (TFORF2860) was obtained from Addgene (plasmid #143643), with ID1 expression driven by the EF-1α promoter. Both constructs were used for lentiviral production and subsequent transduction, as described in previous section.

### Genome characterization

The genetic contents of all hESC were assessed through shallow whole-genome sequencing (performed by BRIGHTcore, UZ Brussels, Belgium), as previously described [52]. The copy number variant analysis targeting gains in 1q, 12p, and 20q11.21 was done using quantitative real-time PCR. Genomic DNA was extracted using the DNeasy Blood & Tissue Kit (Qiagen), quantified via NanoDrop 1000 (Thermo Fisher), and stored at 4°C. For qPCR, reactions (20 µL total volume) contained 10 µL TaqMan 2× MasterMix Plus–Low ROX (Eurogentec), 1 µL TaqMan Copy Number Assay (ID1 [20q], KIF14 [1q], and NANOG [12p]; Table S2), 1 µL RNaseP reference assay (Applied Biosystems), 40 ng DNA, and nuclease-free water. Cycling conditions were: 2 min at 50°C; 40 cycles of 95°C for 10 min, 95°C for 15 sec, and 60°C for 1 min. hPSC DNA with known karyotype and non-template controls were included, with samples run in triplicates on a ViiA7 system (Thermo Fisher). Data was analyzed using Quantstudio Real-Time PCR System v1.3. The assay details are listed in Table S2 and the detailed protocol is available at https://osf.io/jcen3/. All the samples at the initiation of differentiation and the end of neuroectoderm differentiation were verified by qPCR to be free of common chromosomal abnormalities (1q, 12p, and 20q).

### Flow cytometry

Fluorescently labelled hESC^20q^ were dissociated into single cells using TrypLE™ (Thermo Fisher Scientific) for 10 minutes at 37°C. The cell suspension was gently triturated with 2 mL of NutriStem medium and filtered through a 20 µm pluriStrainer to remove cell aggregates. Following centrifugation (1000 rpm, 5 minutes), the cell pellet was resuspended in fresh NutriStem medium. Fluorescent populations were isolated to >95% purity using a BD FACS Aria III (BD Biosciences).

### Neuroectoderm specification

Neuroectoderm differentiation was performed following an adapted protocol [53]. hESC were plated at 122,500 cells/cm² on Laminin-521(5 μg/mL)-coated plates in NutriStem hESC XF medium supplemented with 1:100 RevitaCell (Thermo Fisher). Upon reaching 90% confluence (typically within 24 hours), differentiation was initiated using neural induction medium consisting of DMEM/F-12 (Thermo Fisher) supplemented with 1x NEAA (TFS), 1x GlutaMAX (TFS), 1x 2-mercaptoethanol (TFS), 25 µg/ml insulin (Sigma-Aldrich) and 1x penicillin/streptomycin (TFS). The medium was further supplemented with 10 μM SB431542 (Tocris), 250 nM LDN193189 (STEMCELL Technologies), and 100 nM retinoic acid (Sigma-Aldrich). The differentiation medium was replaced daily for 8 days with supplements.

### Retinal pigment epithelium differentiation

RPE differentiation was performed following modified Plaza Reyes et al. protocol [54–56]. hESC (100,000 cells/cm²) were plated on laminin-521 (10 μg/mL)-coated dishes in NutriStem hESC XF medium with 1:100 RevitaCell (first 24h). At 90% confluence, cells were transitioned to NutriStem hPSC XF GF-free medium (lacking bFGF and TGF-β), with daily medium changes maintained throughout differentiation. The cultures were not replated during the differentiation process, with samples collected at specified time points.

### Total RNA isolation, cDNA synthesis and quantitative real-time PCR for gene expression analysis

Total RNA was extracted from cell pellets using RNeasy Mini/Micro kits (Qiagen) according to the manufacturer’s protocol. RNA concentration was quantified using a NanoDrop 1000 spectrophotometer (Thermo Fisher), and samples were stored at-80°C. For cDNA synthesis, a minimum of 500 ng RNA was reverse-transcribed using High-Capacity RNA-to-cDNA Kit (Thermo Fisher), following the manufacturer’s instructions. The cDNA was stored at-20°C.

Quantitative real-time PCR was performed on a ViiA 7 system (Applied Biosystems) in 20 µL reactions containing: 10 µl of 2x qPCR Master Mix Low ROX (Eurogentec), 1 µl of 20x TaqMan Gene Expression Assay (Life Technologies), 1 µl of nuclease-free water and 40 ng of cDNA. All samples were run in triplicate using standard cycling parameters. Data were analyzed using Quantstudio Real-Time PCR System v1.3 and Gene expression (2^-ΔCt) was normalized to house-keeping gene GUSB (Applied Biosystems). The related TaqMan assay IDs are listed in Table S2.

### Immunostaining

Differentiated cells were washed 3× with PBS and then fixed with 3.6% paraformaldehyde (15 min, RT). After PBS washes (3×), cells were permeabilized with 0.1% Triton X-100 (10 min, RT) and blocked with 10% fetal bovine serum FBS (1 h, RT). Primary antibodies (diluted in 10% FBS) were incubated overnight at 4°C. Following day, cells were washed (PBS, 3×, 5 min) and incubated with Alexa Fluor-conjugated secondary antibodies (1:200 in 10% FBS) and Hoechst 33342 (1:2000) for 1-2 h (RT, dark). After final PBS washes (3×, 5 min), samples were stored at 4°C protected from light. Imaging was performed using a Zeiss LSM800 confocal microscope. The lists with antibodies can be found in supplementary Table S3.

### Microscopy image analysis pipelines

Fluorescence microscopy images from 20q and WT cells were analysed using a custom image-analysis pipeline (IA-20q-REGE-2D). Multi-channel images were segmented at single-cell resolution using custom-trained Cellpose models for cytoplasmic and nuclear detection. For each segmented cell, nuclear area and marker-specific fluorescence intensities were quantified, generating per-cell datasets across experimental replicates. Marker co-localisation within nuclei was used to classify cells into phenotypic categories based on predefined overlap thresholds, and results were summarised using aggregated and per-image visualisations. All analyses were performed in a containerised and version-controlled environment to ensure full reproducibility. All code and configuration files used in this study are available in our GitHub repository to ensure full transparency and reproducibility. The repository contains the complete analysis pipeline, including python scripts and HPC job submission scripts used for segmentation, quantification, and phenotype classification. The repository is publicly accessible at: https://github.com/LIVR-VUB/IA-20q-REGE-2D The custom-trained Cellpose models for whole-cell and nuclear segmentation generated for this study are publicly available via the Hugging Face page for LIVR-VUB. These models were used for all segmentation analyses presented in this work and can be accessed at: https://huggingface.co/LIVR-VUB/20q-REGE-2D

### Single-cell RNA sequencing and data analysis

Differentiated RPE cells (2 months) were gently washed 3× with PBS to remove non-adherent cells. Cell dissociation was achieved using activated Papain solution (Worthington) incubated at 37°C for 30-60 minutes according to the manufacturer’s specifications. For differentiated neuroectoderm cells, single-cell suspensions were achieved using TrypLE™ (Thermo Fisher) treatment (10 min, 37°C), followed by enzymatic neutralization with complete medium and centrifugation at 1000 rpm for 5 minutes to obtain cell pellets.

Approximately 4 million cells were fixed following the manufacturer’s protocol (Parse Biosciences). Post-fixation, cell concentrations were quantified and aliquots stored at −80°C. Single-cell libraries were prepared using the Single Cell Whole Transcriptome Kit v2 (Parse Biosciences), with barcoding, cDNA synthesis, and library generation performed according to the manufacturer specifications. Final libraries were sequenced on an Illumina NovaSeq platform, targeting 20,000 reads per cell.

Raw FASTQ files were aligned to the GRCh38 human reference genome (refdata-gex-GRCh38-2020-A) using the Parse Biosciences pipeline (v0.9.6) with default parameters. Processed gene–cell count matrices were imported into the Seurat R package (v5.3.0) for downstream analysis. Data were log-normalized and highly variable features were identified prior to integration. Integration of multiple sequencing runs was performed using the FindIntegrationAnchors and IntegrateData functions. Dimensionality reduction was carried out using principal component analysis (RunPCA), and Uniform Manifold Approximation and Projection (UMAP) was computed using the first 30 principal components (dims 1–30). Cell clustering was performed using the FindNeighbors and FindClusters functions with a resolution of 1. Visualization was performed using the DimPlot, FeaturePlot, VlnPlot, and DoHeatmap functions. Cell-level gene signature scoring was conducted using the UCell R package, and bubble plots were generated using ggplot2.

### Statistics

All differentiation experiments were performed in at least triplicate (n ≥ 3). Data are presented as mean ± standard error of the mean (SEM) or standard deviation (SD), as indicated. Statistical comparisons between two groups were conducted using unpaired two-tailed Student’s t-tests, while multiple-group comparisons were assessed by one-way or two-way ANOVA, as appropriate. All analyses were performed in GraphPad Prism 9, with p < 0.05 considered statistically significant.

## Acknowledgments

Y.L. was a predoctoral fellow supported by the China Scholarship Council (CSC, grant number 202007650025), C.J., N.K. were predoctoral fellows supported by the Fonds voor Wetenschappelijk Onderzoek Vlaanderen (FWO, grant numbers FWOTM1140 and FWOTM1016 respectively). M.C.D. is a predoctoral fellow supported by the 175 Military Hospital in Vietnam. This research was supported by the FWO (grant number G0713222N) and the Methusalem Grant to Karen Sermon and Claudia Spits (Vrije Universiteit Brussel). The resources and services used in this work were provided by the VSC (Flemish Supercomputer Center), funded by the Research Foundation - Flanders (FWO) and the Flemish Government.

The authors thank Kristoffer Riecken (University of Hamburg) for kindly sharing lentiviral constructs for the expression of the fluorescent proteins mCherry and Venus. We are grateful to An Verloes for proofreading the manuscript and for her support as lab manager. We thank Brecht Ghesquiere for confocal imaging support and Reihaneh Asadi for performing part of the experiments as a master’s student.

## Author contributions

Y.L. and N.K. carried out all the experiments and bioinformatics analysis unless stated otherwise and co-wrote the manuscript. A.S. performed all image quantifications. M.C.D., A.H. and C.J. assisted with DNA/RNA extraction, cell culture and molecular analysis. S.V. assisted with the flow cytometry. Flow cytometry experiments were performed using equipment from the research group of L.A.vG., who also supervised image quantification. D.A.D. and C.S. co-wrote the manuscript and designed and supervised the experimental work.

## Declaration of interests

The Authors declare no Competing Financial or Non-Financial Interests.

## Notes

https://osf.io/8p9qv

### Competing Interest Statement

The authors have declared no competing interest.

