## Supplementary file for "Increases in *BCL2L1* and *ID1* dosage synergistically drive fate bias and competitive advantage in human pluripotent stem cells"

**Supplementary Data**

**Table S1. Karyotype and breakpoints of the hESC lines in this study determined with shallow whole genome sequencing**

| Cell line | Karyotype | Breakpoints in GRCh37/hg19 build | Size |
| --- | --- | --- | --- |
| VUB02^wt^ | 46, XY | NA | NA |
| VUB03^wt^ | 46, XX | NA | NA |
| VUB14^wt^ | 46, XX | NA | NA |
| VUB02^20q11.21^ | 46, XY, dup(20)(q11.21) | chr20:29,650,001-31,100,001 | 1.45 Mb |
| VUB03^20q11.21^ | 46, XX, dup(20)(q11.21) | chr20:29,675,001-33,825,001 | 4.15 Mb |
| VUB14^20q11.21^ | 46, XX, dup(20)(q11.21) | chr20:29,700,001-31,375,001 | 1.68 Mb |

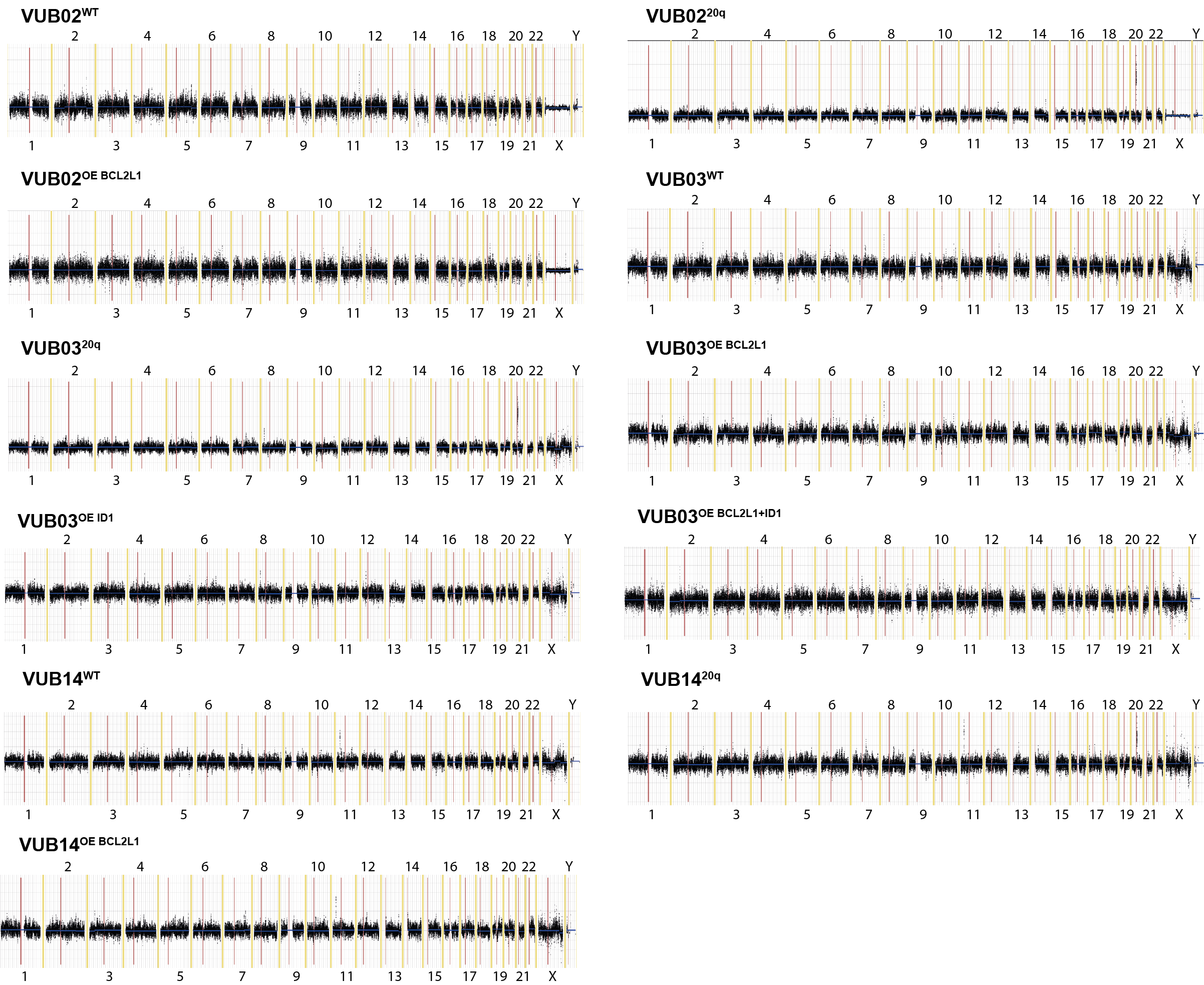

**Figure S1. Shallow DNA sequencing–based karyotyping of the hESC lines used in this study.**

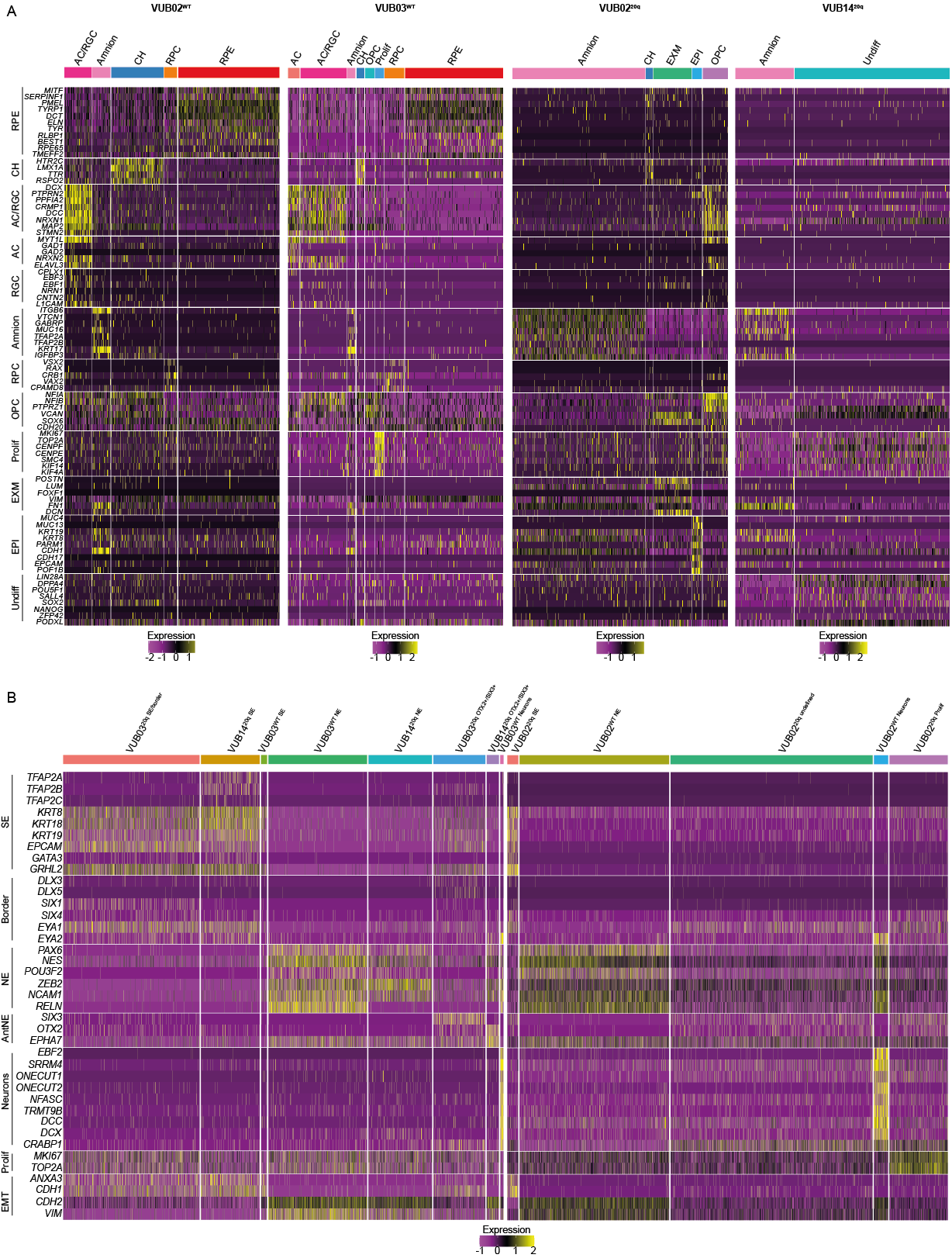

**Figure S2. Heatmap showing the expression of cell-type marker genes across clusters identified in scRNA-seq data from cell lines differentiated toward RPE (A) and neuroectoderm (B). The same gene sets were used for cluster identification in Figure 3A.**

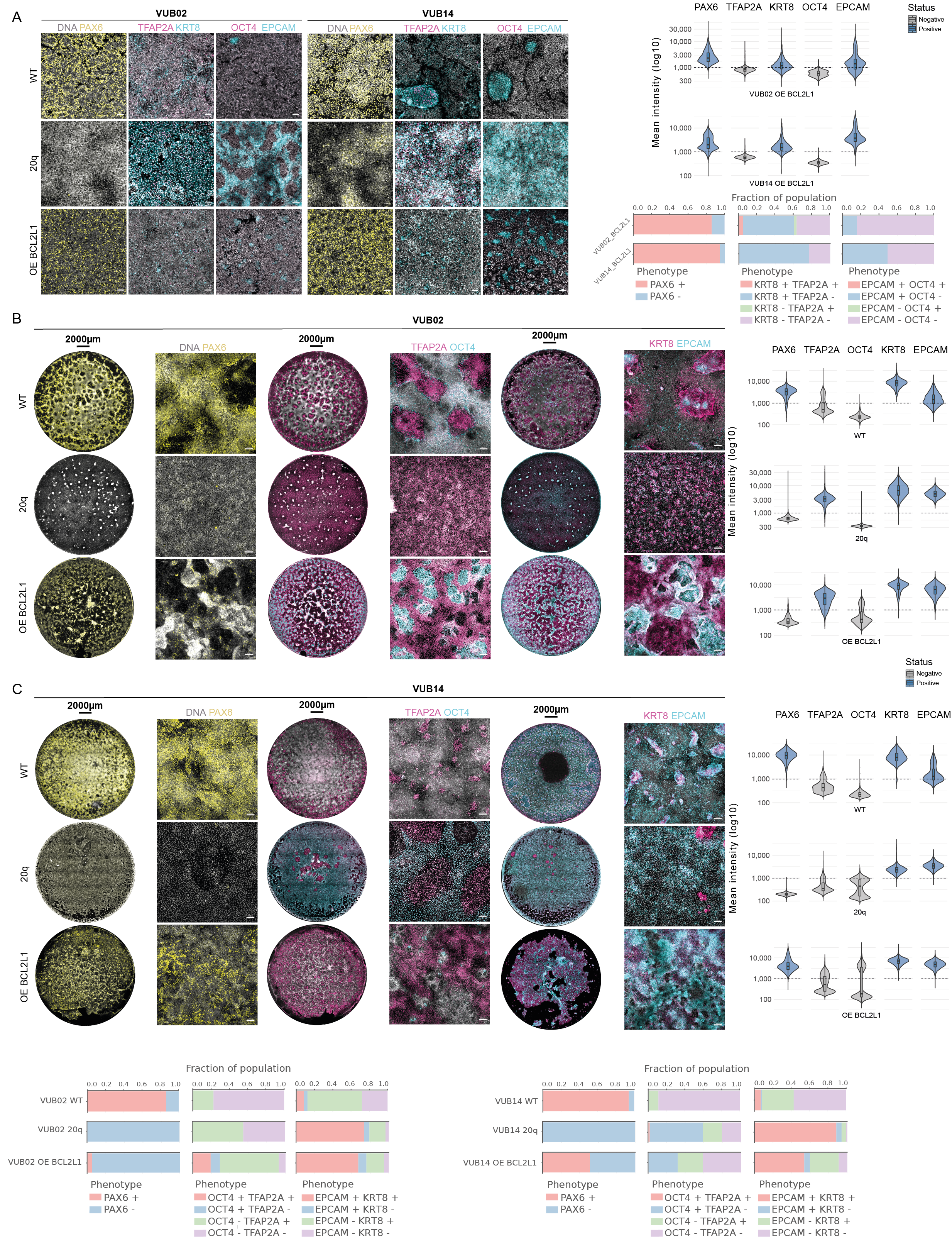

**Figure S3. Immunostaining after differentiation and quantification of marker-positive cells.**

(A) Neuroectoderm differentiation of cell lines VUB02 and VUB14, with quantification of mean fluorescence intensity and the fraction of marker-positive cells.
(B) Differentiation toward RPE for 14 days in cell lines VUB02 and VUB14, with quantification of mean fluorescence intensity and the fraction of marker-positive cells.

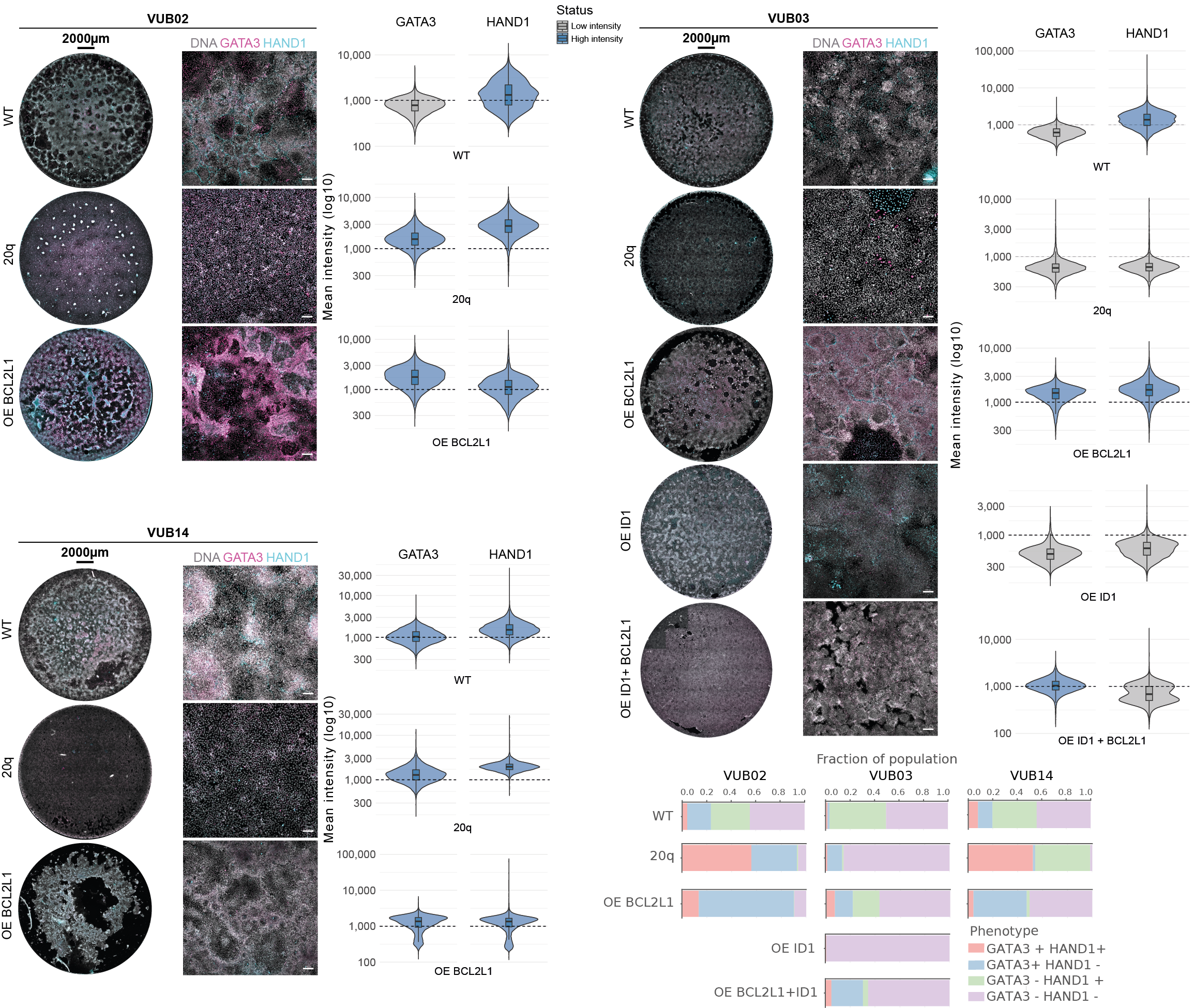

**Figure S4. Immunostaining for the extraembryonic markers GATA3 and HAND1 after 14 days of RPE differentiation, with quantification of mean fluorescence intensity and the fraction of marker-positive cells.**

**Table S2. Gene expression and copy number Taqman assays used in this study.**

Gene expression assays

| Gene | Supplier | Catalog number |
| --- | --- | --- |
| GUSB | Applied Biosystems | 4326320E |
| PAX6 | Thermo Scientific | Hs00240871 |
| SOX1 | Thermo Scientific | Hs01057642 |
| POU5F1 | Thermo Scientific | Hs00742896 |
| BEST1 | Thermo Scientific | Hs00188249 |
| RPE65 | Thermo Scientific | Hs00165642 |
| MITF | Thermo Scientific | Hs01117294 |

Copy number assays

| Gene | Supplier | Catalog number |
| --- | --- | --- |
| RNaseP | Thermo Scientific | 4403326 |
| ID1 | Thermo Scientific | Hs01892845 |
| KIF14 | Thermo Scientific | Hs00637799 |
| NANOG | Thermo Scientific | Hs03820140 |

**Table S3. Antibodies used in this study.**

| Protein | Species | Supplier | Catalog number | Dilution |
| --- | --- | --- | --- | --- |
| PAX6 | Mouse | Abcam | ab78545 | 1:200 |
| OCT4 | Rabbit | Cell Signaling Technology | 2840S | 1:400 |
| TFAP2A | Mouse | Thermo Scientific | MA1-872 | 1:200 |
| PMEL  KRT8  EPCAM  GATA3  HAND1 | Mouse  Rat  Mouse  Rabbit  Goat | Invitrogen  Merck  Thermo Scientific  Abcam  R&D Systems | MA1-34759  MABT329M  14-9326-82  ab199428  AF3168 | 1:200  1:200  1:100  1:250  1:200 |

| Species | Conjugate | Supplier | Catalog number | Dilution |
| --- | --- | --- | --- | --- |
| Donkey anti Mouse | Alexa Fluor 488 | Thermo Fischer Scientific | A-21202 | 1:200 |
| Donkey anti Mouse | Alexa Fluor 594 | Thermo Fischer Scientific | A-21203 | 1:200 |
| Donkey anti Rabbit | Alexa Fluor 488 | Thermo Fischer Scientific | A-21206 | 1:200 |
| Donkey anti Rabbit | Alexa Fluor 594 | Thermo Fischer Scientific | A-21207 | 1:200 |
| Donkey anti Rabbit | Alexa Fluor 647 | Thermo Fischer Scientific | A-31573 | 1:200 |
| Donkey anti Mouse Donkey anti Rat  Donkey anti Rat  Donkey anti Goat | Alexa Fluor 647  Alexa Fluor 488  Alexa Fluor 594  Alexa Fluor 647 | Thermo Fischer Scientific  Thermo Fischer Scientific  Thermo Fischer Scientific  Thermo Fischer Scientific | A-31571  A-21208  A-21209  A-21447 | 1:200  1:200  1:200  1:200 |
| Hoechst 33342 |  | Invitrogen | H3570 | 1:2000 |
